# A Russian validation study of the Coma Recovery Scale-Revised (CRS-R)

**DOI:** 10.1101/276881

**Authors:** Elizaveta G. Mochalova, Liudmila A. Legostaeva, Alexey A. Zimin, Dmitry V. Sergeev, Maxim A. Domashenko, Vladislav Y. Samorukov, Dzhamilya G. Yusupova, Julia V. Ryabinkina, Natalia A. Suponeva, Michael A. Piradov, Yelena. G. Bodien, Joseph T. Giacino

## Abstract

**Introduction**: The aim of the present study was to validate a Russian adaptation of the Coma Recovery Scale–Revised (CRS-R).

**Subjects and methods**: We evaluated 58 patients diagnosed with chronic disorders of consciousness (>4 weeks post-injury, DOC) of various etiology and two patients in a locked-in state at different time points in their post-comatose recovery. We tested sensitivity for changes over 1 week, reliability, as well as criterion validity and diagnostic sensitivity of the Russian adaptation of the CRS-R in comparison with the Russian adaptations of Full Outline of UnResponsiveness Score (FOUR), and Glasgow Coma Scale (GCS).

**Results**: We obtained good sensitivity for changes in neurological status over one week (p<0.0001) and good test-retest reliability (r=1, p<0.0001) of the CRS-R. Inter-rater reliability for the CRS-R total score (κ=0.99, p<0.001) and subscale scores was good. We showed high internal consistency (α=0.87 and 0.89 for the first and second visit respectively). We also showed good criterion validity between two other standardized behavioral scales (moderate correlation with GCS, r=0.597 and high correlation with FOUR Score, r=0.900). CRS-R also demonstrated a significantly higher sensitivity in differential diagnosis of DOC, as compared to GCS, and FOUR Score (p<0.001).

**Conclusion**: The results show that the Russian version of the CRS-R is a valid and sensitive tool for the evaluation of severely brain damaged patients with chronic DOC which can be used for differential diagnosis and for the assessment of dynamic recovery.

## INTRODUCTION

Progress in critical care has led to an increase in the number of patients who survive severe acute head trauma and non-traumatic brain injury. This has resulted in a rising number of patients being diagnosed with chronic disorders of consciousness (DOC), such as vegetative state (VS), also known as unresponsive wakefulness state (i.e. demonstrating spontaneous eye-opening without signs of awareness) [1], and minimally conscious state (MCS, evidence of inconsistent but clear and reproducible signs of awareness) [2]. Objective evaluation and distinguishing between different chronic DOC states is of paramount importance for diagnosis, prognosis, and determining rehabilitation potential: patients with MCS have better long-term outcome [3, 4]. However, accurate differentiation between various types of chronic DOC may be challenging, as subtle behavioral signs of consciousness can easily be missed. In this context, various standardized diagnostic clinical tools were developed. The Coma Recovery Scale — Revised (CRS-R) [5] is recognized as the most comprehensive and reliable behavioral assessment for DOC [5-7]. In a review by the Archives of Medicine and Rehabilitation, it was the only tool out of 13 other consciousness scales that was recommended for assessment of DOC with only minor reservations [6]. The CRS-R has been translated into several languages with validation studies completed in Spanish, Italian and French [8-10].

Several scales for chronic DOC assessment were developed in Russia, including Dobrokhotova-Zaytsev scale [11] and communicative activities scale designed by Bykova and Lukianova for pediatric practice [12]. They are not available in other languages. Unfortunately, wide use of internationally approved specific scales for chronic DOC evaluation is limited in Russia due to a lack of translated and validated versions.

The Glasgow Coma Scale (GCS) [13] appears to be the most widely used tool for assessing DOC in the Russian population, although a Russian translation of the scale has never been validated. The GCS was developed to monitor severely brain injured patients in the acute care setting and does not allow for differential diagnostic assessment in suspected chronic DOC [6, 14]. In the context of chronic DOC examination, GCS does not specifically test for MCS by Aspen criteria [2] and does not account for intubated patients. There is also poor reliability, prognostic validity for DOC [9], and GCS is not standardized. There is no assessment of visual fixation and pursuit, object recognition and intentional communication, behaviors that are very important for diagnosing MCS. In fact, the only items to assess MCS patients are found on the motor response subscale (i.e., localization to pain, command following) and verbal response assessment (confused verbal response, inappropriate words). Another scale that is used in DOC, mainly in the acute period, is the Full Outline of Unresponsiveness Score (FOUR)[15] In contrast to the GCS, the FOUR was validated and demonstrated greater inter-rater reliability than the GCS in patients in the emergency department [15-17]. In contrast to the GCS, the FOUR Score assesses visual pursuit and visual fixation which are among the most sensitive behaviors for differentiation of MCS and VS [18]. Additionally, the FOUR Score may be used in chronic patients [19]. But still FOUR is less informative for assessing chronic DOC than CRS-R and is not adequate for differential diagnosis between VS and MCS [19].

Absence of validated Russian-language scales for assessment of DOC significantly hinders both clinical and scientific research activities involving this category of patients. First, absence of a tool for sensitive detection of consciousness is a notable problem in rehabilitation and clinical prognosis determination. Second, clinical results and research data from different sites cannot be juxtaposed or consolidated, which, given the typically small size of this patient population, may significantly reduce the statistical power of investigations. One solution to ensuring more accurate assessment of DOC and providing a platform for data sharing is to validate a translated Russian version of the CRS-R. A validated Russian version of the CRS-R will provide a common tool for monitoring chronic DOC patients in all Russian-speaking territories. The aim of this investigation is to assess the validity, reliability, diagnostic sensitivity of the Russian version of the CRS-R and to determine its ability to monitor changes in neurobeahavioral status over time.

## MATERIALS AND METHODS

### CRS-R TRANSLATION AND CULTURAL ADAPTATION

In the first stage of this study, we completed the following steps to translate the CRS-R from English into Russian:

1. Two medical translators independently translated the CRS-R into Russian.
2. We combined the two translations into a single scale.
3. A native English speaker fluent in Russian, with a medical education, translated the Russian CRS-R back into English. This translator was not involved into the translation of the scale into Russian.
4. The author of the original CRS-R and a Russian-speaking collaborator evaluated the translation and back-translation and provided suggestions for modification of the translation. We revised the Russian scale to ensure concordance between measures.
5. We obtained final approval of the back translation from the original author of the CRS-R.

Full Russian CRS-R is available in Supplement 1.

### VALIDATION OF THE RUSSIAN CRS-R SCALE

In the prospective validation phase of this study, we enrolled patients from a specialized DOC program and acute medical centers who met the following inclusion criteria: (1) age 18 years and older and (2) clinically defined DOC, including emerging MCS (eMCS), or locked-in state (LIS) (3) at least 4 weeks aftertraumatic or non-traumatic brain injury resulting in coma. LIS is a condition that should be considered when making a differential diagnosis in patients with acute or chronic disorders of consciousness [22], and in these cases CRS-R may be helpful. Therefore, we included these patients in our study to determine concordance between the Russian and English version of CRS-R assessment of this rarely occurring cohort. We excluded subjects who were diagnosed with coma, including irreversible coma/brain death, received sedatives or central muscle relaxants at the time of clinical evaluation, had injury to both eyes or tympanic membranes, or inner ear, severe trauma to both upper or lower extremities, or unstable physical condition.

The study was approved by the local ethics committee and registered at clinicaltrials.gov (Identifier IDNCT03060317). During the evaluation period, all patents continued to receive standard medical treatment and rehabilitation.

Three neurologists with at least 3 years’ experience in management of chronic DOC patients independently evaluated participants with the Russian adaptation of the CRS-R. The 3 examiners watched a training video and reviewed the administration and scoring guidelines before using the CRS-R.

The validation procedure followed the method employed for validation of the original CRS-R scale [5]. Two independent examiners (A and B) administered the CRS-R on Day 1. Examiners A and B remained blind to each other’s evaluations but knew the clinical medical diagnosis of the patients. Examiners evaluated participants while they were in a wakeful state, with their eyes open, and the CRS-R arousal facilitation protocol was applied when necessary during the assessment. There was a limit of 5 applications of the arousal facilitation protocol to assure the arousal of the patient. During the Day 1 visit, both experts assessed patients consecutively (evaluations A1 and then B1). Then examiner A re-assessed all patients on the same day (evaluation A1-retest). The CRS-R was administered again at the second visit (7 days later) by the same examiners (evaluations A2 and B2) to investigate the sensitivity of the CRS-R to change over time.

Reliability shows that the scale produces similar results under consistent conditions, when used by different investigators (inter-rater) or in different examinations (retest) [20]. Assessment of reliability included internal consistency (i.e., total scores and subscales of A1 and B1), inter-rater reliability (i.e. total scores of A1 and B1) and test-retest reliability (i.e. the CRS-R total score obtained between A1 and A1-retest evaluations).

Criterion validity is the extent to which a measure is related to an outcome, which is assessed by calculating the correlation between a new scale and a customary or validated one [20]. To assess CRS-R criterion validity, we administered two behavioral scales with established reliability, the GCS [13] and the FOUR Score [15]. Neither the GCS nor the FOUR Score are validated in Russia, but both have been translated into Russian and are commonly used in clinical practice. Both scales were administered by examiners A and B on Day 1 after CRS-R assessment.

The diagnostic sensitivity of CRS-R was tested via comparison of the MCS diagnosis established with the following three scales: the CRS-R, the GCS and the FOUR, all of which were administered during the first visit (A1 and B1). So the rater administering the GCS/FOUR was not blind to the CRS-R assessment. We assigned diagnostic categories to the behaviors assessed on both scales based on the Aspen criteria [2] (as described in the French CRS-R validation study (Table 1) [9].

**Table 1.** Criteria for establishing MCS using CRS-R, GCS and FOUR (sources: [2, 9]).

| Scale | MCS criteria |
| --- | --- |
| CRS-R | One of the following:<br>Auditory function: scores 3-4 (consistent movement to command (Au4), reproducible movement to command (Au3))<br>Visual function: scores 2-5 (object recognition (V5), object localization (V4), visual pursuit (V3) and fixation (V2))<br>Motor function: 3-6 scores (functional object use (M6), automatic motor response (M5), object manipulation (M4) and localization to noxious stimulation (M3))<br>Verbal function: score 3 (intelligible verbalization, Ver3)<br>Communication: scores 1-2 (functional (C2) and non-functional (C1) communication) |
| GCS | One of the following:<br>Verbal responses: scores 3-4 (confused verbal response (V4), inappropriate words (V3))<br>Motor responses: scores 5-6 (obeying commands (M6), localizing pain (M5)) |
| FOUR | One of the following:<br>Eye responses: score 4 (tracking, or blinking to command, E4)<br>Motor responses: scores 3-4 (thumbs up, fist or peace sign (M4); localizing to pain (M3)) |

### STATISTICAL ANALYSIS

We present descriptive data on patient characteristics as means and standard deviations in the case of normally-distributed data and as medians and interquartile intervals for non-normal distributions. We used the Mann–Whitney U test to establish any difference between the evaluations of patients by two experts using CRS-R on Day 1.

We used the non-parametric Wilcoxon signed-rank test to compare the A1 (Day 1) and A2 (Day 2, 7 days later) scores in order to test CRS-R sensitivity to changes in level of consciousness over a one week period [5]. All tests employed level of significance α = 0.05.

We used Cronbach’s alpha to assess the reliability, or internal consistency of the scale. We compared We considered the value of 0.8 as a reasonable threshold indicating “good” reliability [10].

We used **Cohen’s Kappa coefficient and inter-class** correlation to measure inter-rater agreement between the overall CRS-R scores, and between the scores per individual test item given by the two experts [9]

We assessed the test-retest reliability/repeatability with Pearson correlation coefficient. We applied Spearman’s rank correlation coefficient to measure the degree of values association between two ranked variables (single or multiple) from different scales [10].

The interpretations of strength of correlation is presented in the table 2 [21].

**Table 2.** Rule of Thumb for Interpreting the Size of a Correlation Coefficient [21].

| Size of Correlation | Interpretation |
| --- | --- |
| .90 to 1.00 (-.90 to -1.00) | Very high positive (negative) correlation |
| .70 to .90 (-.70 to -.90) | High positive (negative) correlation |
| .50 to .70 (-.50 to -.70) | Moderate positive (negative) correlation |
| .30 to .50 (-.30 to -.50) | Low positive (negative) correlation |
| .00 to .30 (.00 to -.30) | Little if any correlation |

The capacity of the CRS-R and the two other coma scales (GCS and FOUR) were explored to identify patients in MCS, relative to VS. Chi-squared analyses were performed to compare the proportion of MCS diagnoses between scales and the proportion of MCS diagnosis. For estimation of differential diagnostic sensitivity of the CRS-R, we divided the patients into the groups of VS and MCS using criteria for CRS-R, FOUR and GCS separately (Table 1). We measured the significance of the difference in proportion between the scores in these two groups with the non-parametric Wilcoxon signed-rank test.

We used IBM SPSS Statistics 22 package for statistical analysis [22].

## RESULTS

The study was conducted between October 2016 and April 2017. Participants were 58 patients with chronic DOC and 2 patients with LIS of various etiology and duration admitted to 4 clinical hospitals and research centers in Moscow. Age ranged between 18 and 84 years, 31 patients were male. Sample characteristics are shown in the Table 3.

**Table 3.** Sample characteristics.

| Category | Number |
| --- | --- |
| Number of subjects | 60 |
| Age, yr (mean±standard deviation) | 46±18 |
| Time post-injury, months (median [interquartile range]) | 2.5 [1;7] |
| Gender distribution, male/female | 31/29 |
| Etiology: |  |
| - Trauma | 8 |
| - Anoxia | 22 |
| - Stroke | 25 |
| - Demyelinating disease | 1 |
| - Infection meningitis/encephalitis | 2 |
| - Metabolic (toxic brain damage, pontine myelinolysis) | 2 |
| Clinical state according to Aspen Criteria: |  |
| - VS | 27 |
| - MCS and eMCS | 31 |
| - Locked-in state | 2 |

### OVERALL ASSESSMENT RESULTS

Patients were evaluated twice within a one-week interval (mean 8.8±2.7 days) to assess the CRS-R’s sensitivity to change in diagnostic status during this time interval. Median CRS-R total score after the first visit A1 was 8.5 [5.0; 14.75] and after the second visit (A2) – 10.0 [5.0; 17.75], p<0.0001. In 11 patients (18,3%), this change was associated with a change in clinical diagnosis, suggesting sufficient sensitivity in capturing changes in diagnosis (Fig. 1). Of note, such changes were documented in early post-coma period, reflecting the natural course of the disease.

**Figure 1.**
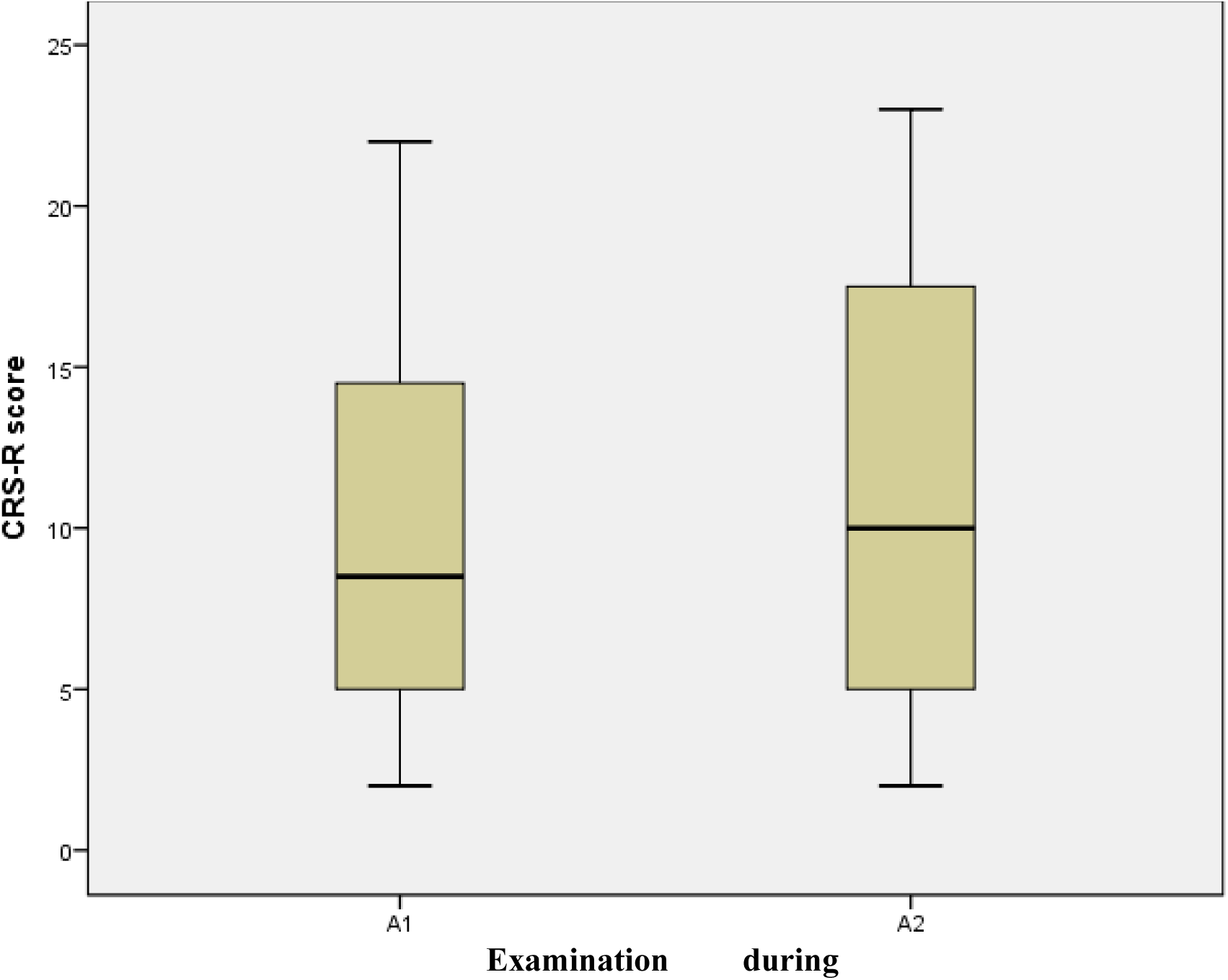
Distribution of CRS-R total scores acquired during a 1-week interval by the first examiner (A1 and A2).

### CRS-R RELIABILITY

Internal consistency of the total score using Cronbach’s alpha was high 0.87 (p=.0001),

Inter-rater reliability was assessed using Cohen’s kappa to determine the reproducibility of CRS-R total score and subscale scores between the different observers (Table 4). The data obtained indicate very close agreement between the two experts. Subscale score agreement was also high. The results are presented in Table 5. demonstrating reproducibility of assessment results by different experts: mean score for A1 10.25±5.80 and for B1 10.20±5.84.

**Table 4.**
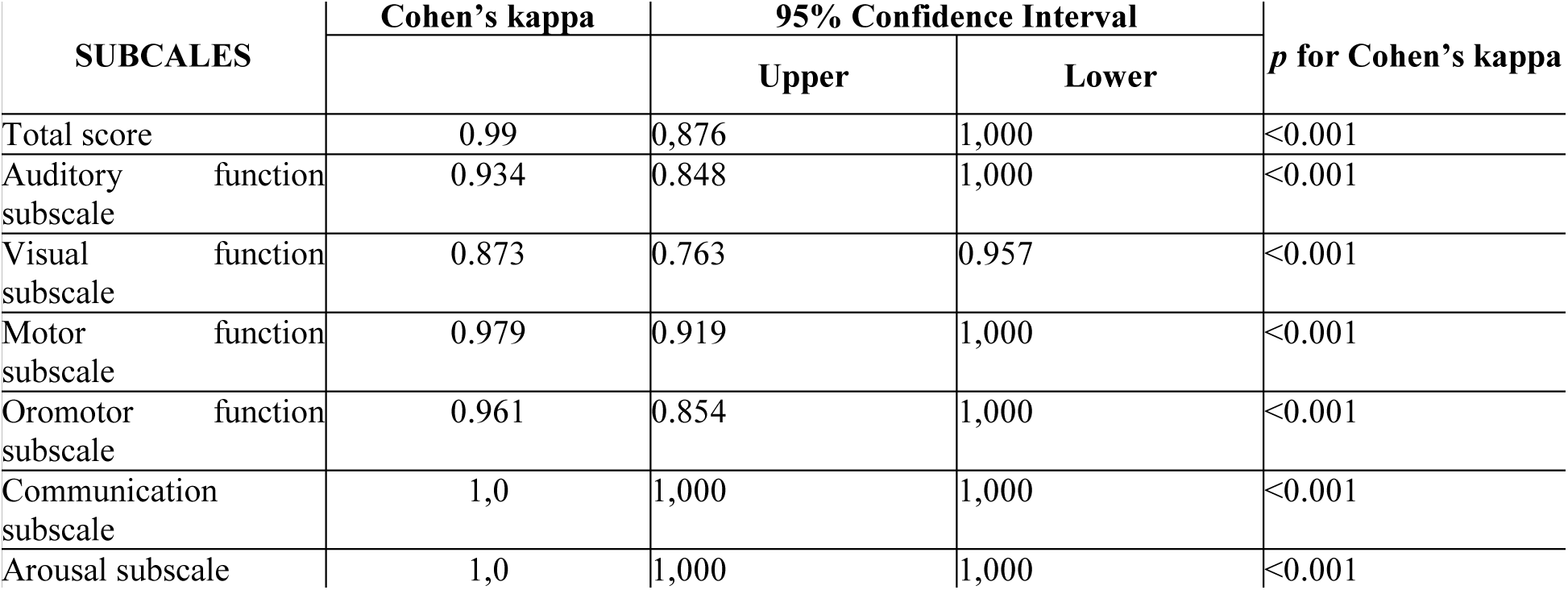
CRS-R reliability assessment via inter-rater assessment.

| SUBCALES | Cohen's kappa | 95% Confidence Interval |  | *p* for Cohen's kappa |
| --- | --- | --- | --- | --- |
|  |  | Upper | Lower |  |
| Total score | 0.99 | 0,876 | 1,000 | <0.001 |
| Auditory function subscale | 0.934 | 0.848 | 1,000 | <0.001 |
| Visual function subscale | 0.873 | 0.763 | 0.957 | <0.001 |
| Motor function subscale | 0.979 | 0.919 | 1,000 | <0.001 |
| Oromotor function subscale | 0.961 | 0.854 | 1,000 | <0.001 |
| Communication subscale | 1,0 | 1,000 | 1,000 | <0.001 |
| Arousal subscale | 1,0 | 1,000 | 1,000 | <0.001 |

**Table 5.** CRS-R criterion validity: correlation between CRS-R and GCS, FOUR total scores and similar subscales.

| Scale and Subscale | R | p |
| --- | --- | --- |
| CRS-R: total score |  |  |
| GCS: total score | 0.90 | 0.0001 |
| FOUR: total score | 0.60 | 0.0001 |
| CRS-R: visual function |  |  |
| GCS: eyes opening | 0.26 | 0.047 |
| FOUR: eye responses | 0.70 | 0.0001 |
| CRS-R: motor function |  |  |
| GCS: motor function | 0.89 | 0.0001 |
| FOUR: motor response | 0.88 | 0.0001 |
| CRS-R: verbal function |  |  |
| GCS: verbal function | 0.699 | 0.0001 |
| CRS-R: communication |  |  |
| GCS: verbal function | 0.691 | 0.0001 |

Test-retest consistency was also high with the correlation coefficient r=1 (p<.0001), indicating the stability of patient’s assessment during the short observation period during Day 1 (A1 and A1-retest evaluations), suggesting that the Russian CRS-R scale is relatively resistant to change associated with the time factor (Fig. 2).

**Figure 2.**
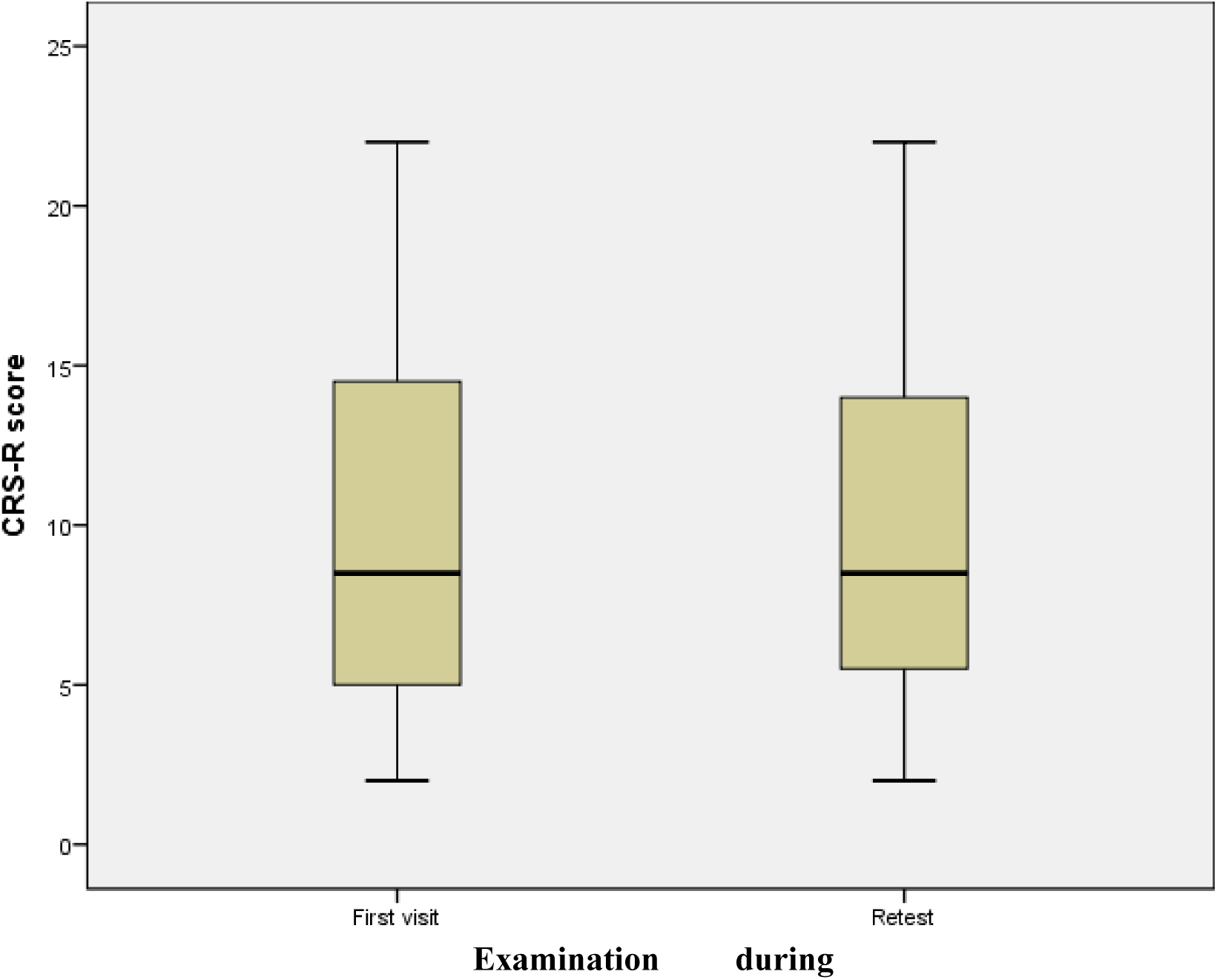
Test-retest distribution of total score CSR-R during Day 1 **(evaluations A1 and A1-retest).**

### VALIDITY ASSESSMENT

Criterion validity was determined by comparing CRS-R total scores to the GCS and the FOUR total scores, as well as comparing their subscale scores. The total CRS-R score showed moderate correlation (r=0.60, p=) with the total FOUR score and very high correlation (r=0.90) with the total GCS score.

We assessed the correlation between CRS-R subscale scores and similar subscales of the FOUR and GCS (Tab. 5). All selected CRS-R subscales showed significant (p<0.05) very high, high or moderate correlation with similar items on the other scales, except for visual function on the GCS (r= 0.257; p<0.05), where the correlation was significant but poor.

### SENSITIVITY FOR DIFFERENTIAL DIAGNOSIS

CRS-R sensitivity for differential diagnosis was assessed in comparison to other scales to estimate the CRS-R’s potential to distinguish between patients in chronic VS and MCS. EMCS and LIS patients were excluded due to the small sample size of these cohorts. It is important to note that both experts were unanimous in the assessment of VS and MCS, so there was no need to compare their results. The total CRS-R score was 5 [4.5; 6.0] in the group of VS patients vs. 13 [10; 19] in the MCS group (p<0.0001). Statistically significant differences were also found between these groups comparing respective CRS-R subscales scores of VS and MCS (p<0.0001)

We assigned each patient a DOC diagnosis using each of the three behavioral tools with particular attention to the MCS diagnosis (Table 6). The capacity of the CRS-R and the two other coma scales (GCS and FOUR) were explored to identify patients in MCS, relative to VS. Chi-squared analysis showed that the proportion of patients diagnosed with MCS by the CRS-R was significantly higher as compared to the GCS (χ^2^=25; p<0.0001) andthe FOUR (χ^2^=36; p<0.0001). In comparison with the CRS-R, 6 patients were misdiagnosed with MCS by FOUR scale, and 11 patients by GCS. Of note, every patient diagnosed with MCS according to the GCS and the FOUR scale met the CRS-R MCS criteria. Visual function, e.g. fixation, pursuit and object recognition, were the most frequent behaviors contributing to the MCS diagnosis.

**Table 6.**
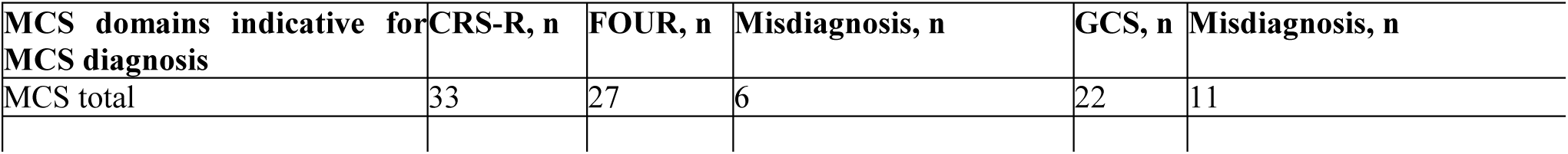
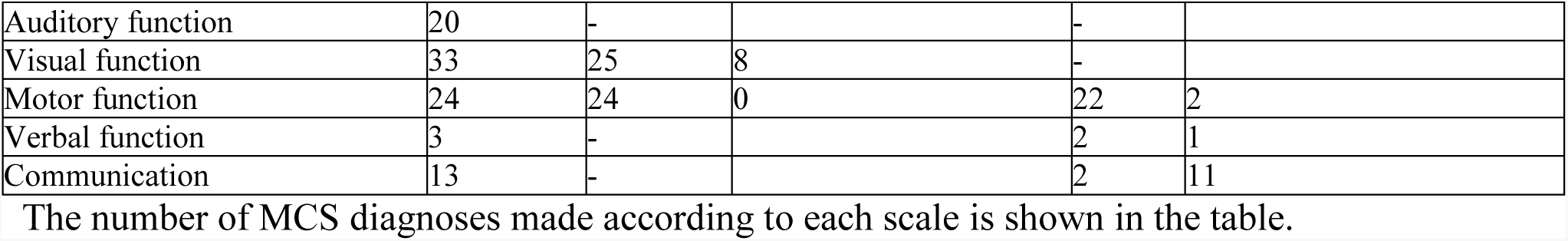
Rates of MCS diagnosis based on total and subscale scores by CRS-R, FOUR and GCS from 60 DOC patients.

Assessment of motor and verbal functions were more prone to erroneous interpretation with the GCS and FOUR scale (Table 7).

**Table 7.** Missed MCS signs in patients diagnosed as MCS using CRS-R when assessed with GCS and FOUR scale.

| Missed MCS signs according to CRS-R | GCS, n | FOUR, n |
| --- | --- | --- |
| Visual fixation | 7 | 6 |
| Visual pursuit | 2 | - |
| Object recognition | 5 | - |
| Object manipulation | 1 | 1 |
| Intelligible verbalization | 1 | - |

The comparison of VS and MCS groups shows that the CRS-R has significant difference in total score and every subscale score (see Table 8).

**Table 8.** Assessment of diagnostic sensitivity: CRS-R, FOUR and GCS scales sensitivity for the differential diagnosis of chronic DOC.

| Scale: Diagnosis, n | Total | Auditory | Visual | Motor | Verbal function | Communication/speech | Arousal | Brainstem reflexes | Respiratory pattern |
| --- | --- | --- | --- | --- | --- | --- | --- | --- | --- |
| **CRS-R: VS, 27** | 5 [4.5;6] | 1 [1;1] | 0 [0;1] | 1 [1;2] | 1 [1;1] | 0 [0;0] | 2 [2;2] |  |  |
| **CRS-R: MCS, 33** | 13 [10;19] | 3 [2;4] | 3 [2;5] | 3 [2;5] | 1 [1;2] | 1 [0;1] | 2 [2;3] |  |  |
| **p (VS, MCS)** | 0.0001 | 0.0001 | 0.0001 | 0.0001 | 0.0001 | 0.0001 | 0.0001 |  |  |
| **FOUR: VS, 33** | 12 [8;13] |  | 3 [3;3] | 2 [1;2] |  |  |  | 4 [3;4] | 4 [0;4] |
| **FOUR: MCS, 27** | 14 [12;16] |  | 4 [4;4] | 4 [3;4] |  |  |  | 4 [4;4] | 4 [1;4] |
| **p (VS, MCS)** | 0.0001 |  | 0.0001 | 0.0001 |  |  |  | **0.017** | **0.838** |
| **GCS: VS, 38** | 8 [7;9] |  | 4 [4;4] | 3 [2;4] |  | 1 [1;1] |  |  |  |
| **GCS: MCS, 22** | 11 [11;12] |  | 4 [4;4] | 6 [6;6] |  | 1 [1;2] |  |  |  |
| **p (VS, MCS)** | 0.0001 |  | **0.859** | 0.0001 |  | 0.0001 |  |  |  |

VS and MCS patients were grouped based on individual scale criteria, respectively (Table. 2). Scores by scales and subscales are presented as Me [LQ; UQ] alongside with the significance of Me differences derived for each scale in Mann-Whitney cross-comparison test in both groups.

At the same time, the FOUR Score showed significant difference in total score and in visual and motor function subscales and brainstem reflexes. The assessment of respiratory pattern did not show statistically significant differences between the groups (p<0.838), probably due to the fact that the spontaneous breathing does not reflect the function of consciousness, and may be affected by extracerebral conditions.

The GCS scale demonstrated significant difference between VS and MCS in total scores, verbal and motor responses (p<0.0001), and failed to show differences in the visual subscale (p<0.859).

Thus, the differences between total scores on all three scales show that each of them can be used for differential diagnosis of VS and MCS. However, the error rates (Table 4) suggest that the assessment with FOUR and GCS scales may miss out MCS cases more often than using the CRS-R (in 10% and 18% of cases, respectively). It should be noted, that every CRS-R domain shows significant difference between VS and MCS patients, and therefore every subscale is informative and assesses an important aspect of the status, unlike the FOUR and GCS, where some items may have low discriminative value.

## DISCUSSION

In the absence of a reliable assessment tool for patients with chronic DOC in Russia, we translated the CRS-R into Russian and tested its validity and reliability. We chose the CRS-R because of its strong psychometric properties [5] and recommendations based on the American Congress of Rehabilitation Medicine [6]. The design of this study was based on CRS-R validation studies performed by our colleagues from other countries [5, 8-10]. The GCS and the FOUR score were selected for comparative testing as these tools appear to be the most familiar to medical specialists in Russia for DOC evaluation, though, these scales are not yet officially translated into Russian. Despite the fact that these two scales are not recommended for the assessment of chronic DOC patients [6] they are concise and easy to administer, and thus became widely used in clinical practice [19] and in CRS-R validation studies [9]. Several other tools for the evaluation of chronic DOC patients are available, such as the Disability Rating Scale (DRS) [23] and the Wessex Head Injury Matrix (WHIM) [24] proved to be more sensitive and applicable to DOC than the GCS and the FOUR, and demonstrating better correlation with the CRS-R scores [9, 10] Unfortunately, these scales are not widely known in Russia, not yet translated and validated, therefore they were not used in our study.

Our sample consisted of adult males and females recovering from coma of different duration. DOC was documented in 58 patients, all of which were more than 4 weeks post-coma. Two patients were diagnosed with LIS. A variety of factors that can interfere with adequate assessment of consciousness were excluded in this study, i.e., use of medications that may cause depression of consciousness, or use of muscle relaxing agents, exogenous and endogenous intoxication, and severely compromised somatic status.

Inclusion of patients with chronic DOC provided an opportunity to evaluate CRS-R sensitivity to changes in patient status within a relatively short period of time. We included 60 patients meeting eligibility criteria. The sample size was selected because of low DOC incidence, capability of enrollment of these patients and the statistical assessment of the sample size. The Aspen criteria were used for differential diagnosis of chronic DOC as the current “gold standard” for detection of MCS [2]. It is shown that GCS and FOUR scales have low sensitivity for MCS detection [6, 14, 19], so we did not rely on them.Different features of the Russian version of the CRS-R were explored in the present study. In particular, the Russian CRS-R adaptation showed high reliability and internal consistency based on the total score and subscales score values. The test-retest interrater reliability procedure yielded high subscale scores indicating close similarity in ratings from two independent observers. The data obtained are consistent with the results from previous CRS-R validation studies, including the original version [5, 8-10].

Content and criterion validity assessments yielded meaningful results. We established a strong relationship between the CRS-R and the GCS, and moderate correlation between the CRS-R and the FOUR. The difference in the strength of the relationship lies in the intrinsic structure of the two comparison scales. Although both scales are used for DOC evaluation, the FOUR envisages assessment of brainstem reflexes and respiratory pattern, usually not indicative of the level of consciousness and varying in patients with chronic DOC, although very important in the assessment of acute DOC.

Assessment of the correlation of individual test items showed satisfactory or strong significant correlations between corresponding items in the CRS-R and other scales, except for discordance in CRS-R visual scale and GCS eyes opening. This discrepancy is also explained by the scales structure: the GCS eye opening test does not imply the assessment of higher cortical visual function, it only helps to estimate the level of patient’s arousal (spontaneous or voluntary eye opening or in response to some stimuli). Despite the difference in total scores, correlations between individual items were higher with the FOUR scale, once again indicating that specific test items on the FOUR do not assess consciousness in chronic DOC patients, likely accounting for discrepancies in total scores between these two scales.

Thus, the analysis of single and multiple variables correlation demonstrate strong and reliable relationship, confirming once again CRS-R high content validity. At the same time high correlation levels demonstrate excellent criterion validity of this scale. Similar data in the assessment of CRS-R criterion validity was obtained in other studies [9, 10].

Inter-rater reliability between the two experts was analyzed. The two raters showed very close and consistent results during re-examination within the short period of time. CRS-R has also been found sensitive to changes in patients’ condition during the follow up, thus, it can be used to monitor the effectiveness of patients’ rehabilitation. Documented changes in DOC patients state are explained by inclusion of several persistent cases in the sample, who managed to achieve some progress in the recovery.

The diagnostic sensitivity of CRS-R for VS and MCS was the final analysis of the study. There was no disagreement between observers in the CRS-R subscale scores for each individual patient. The CRS-R was more sensitive compared to the other scales in MCS identification, although all scales demonstrated significant between-group differences in total scores. The FOUR scale yielded a misdiagnosis in 6 patients, the GCS in 11 patients. Chi-squared analysis showed that the proportion of patients diagnosed with MCS by the CRS-R was significantly higher as compared to the other two scales. These results are similar to our French colleagues’ data [9]. The observed differences may arise from the fact that both scales were designed for assessment of acute DOC, thus, these scales do not account for such symptoms as sustained fixation or visual pursuit (the GCS) or object manipulation and intelligible verbalization (the GCS and the FOUR). In addition, the CRS-R accounts not only for speech production, but also any other form of communication, for example, using gestures or an alphabet board, thus expanding the scale’s potential in accurately establishing the differential diagnosis.

It has been shown that eyes responses such as sustained gaze fixation and visual pursuit appear to be the most common MCS symptoms. These behaviors were identified in all MCS patients from the study sample. Nine patients (15%) did not have any other signs of MCS, which is consistent with published data [4, 10]. It should be mentioned that the FOUR scale does not assess eye fixation, but only visual pursuit, while the GCS does not assess the gaze fixation or pursuit, only eyes opening. This explains the high rate of missed MCS cases.

Only two mistakes were found in motor function evaluation by the GCS, the FOUR results were similar to CRS-R scale values. The FOUR does not assess the verbal function and communication, while the GCS fails to provide adequate evaluation of these functions, thus identifying only 2 MCS patients out of 13 identified by the CRS-R.

Administration of individual subscales identified specific diagnostic features in both groups. Each of CRS-R subscales show statistically significant differences in detection of VS and MCS features, indicating each single item’s high accuracy and sensitivity and absence of non-informative criteria for DOC assessment in the scale structure.

Analysis of the two comparison scales identifies irrelevant items, such as eye opening in the GCS, and respiratory pattern and brainstem reflexes in the FOUR. As previously discussed, the GCS eye-opening item assesses arousal level, and addresses the question, “under what circumstances does a patient open his/her eyes,” It does not assess attentional function (i.e., whether a patient can fix the gaze, pursue or recognize objects etc.), therefore the highest score was given to almost all patients despite differences in diagnosis. Breathing pattern or brainstem reflex assessments are not associated with level of consciousness, but rather reflect the severity of neurological deficit. Such patients may have, for example, brainstem injury, require a ventilatory support, lack some brainstem reflexes, but be able to demonstrate other signs of conscious behavior. These signs are not associated with level of consciousness, supported further by our findings.

The Russian adaptation of CRS-R seems to be the most sensitive scale in terms of VS and MCS differential diagnosis, accurately identifying the minimal signs of awareness and consciousness. The CRS-R is a very useful tool for assessing DOC patients, with prior reports suggesting that it detects 99% of MCS signs [18] and can be administered by people with different skills and education. The CRS-R shows strong psychometric properties [25] and training requirements are minimal [26], though the level of experience may still influence on the results.

The limitations of this study include a modest sample size and a limited number of experts performing clinical evaluation. Nevertheless, power calculation shows that 60 subjects is adequate for a study assessing a 23-item scale. Compared to other CRS-R validation studies, the number of patients in the Russian validation procedure appears sufficient. Prior studies enrolled 35-80 patients. [5, 8-10] Experts participating in the study were experienced neurologists trained in the same center, thus they had comparable expertise, similar approaches to examination of patients, interpretation of signs and CRS-R application, yielding identical results of patients’ evaluation. In this study, the CRS-R was assessed versus the GCS and the FOUR, both of which were designed for use in the acute setting. It should also be noted that the CRS-R, GCS and FOUR assessments were conducted by the same specialists, so. In the present study, only trained neurologists were authorized to use the abovementioned scales

In sum, the study results demonstrate that the Russian adaptation of CRS-R is comparable to other commonly accepted and used scales to assess the DOC, but it has a number of advantages. At present, it is the only validated tool for assessing patients with chronic DOC in Russia.

## CONCLUSION

Findings from this investigation demonstrate that the Russian CRS-R is a valid and reliable tool for assessment of chronic DOC. Standardized behavioral assessments of consciousness are critical for accurate diagnosis, prognosis, and treatment planning of DOC. A validated tool for assessment of DOC in Russian patients will facilitate comparisons of research results across the world and foster collaborations aimed at improving clinical management of this patient population. The Russian CRS-R is currently the only validated standardized tool for assessment of DOC in Russian-speaking countries and should be considered for adoption into clinical practice and research investigations.

## ACKNOWLEDGEMENTS

The study is supported by Russian Science Foundation grant 16-15-00274 (E.M., L.L., J.R., N.S., and M.P.).

The authors highly appreciate efforts of Dr. M. A. Loskutnikov and Dr. V. A. Shandalin in recruiting patients into the study, and are thankful to Sofia Gutkin for her input in the translation of the CRS-R.

## REFERENCES

1. Laureys, S. and N.D. Schiff, Coma and consciousness: paradigms (re)framed by neuroimaging. Neuroimage, 2012. 61(2): p. 478–91.

2. Giacino, J.T., et al., The minimally conscious state: definition and diagnostic criteria. Neurology, 2002. 58(3): p. 349–53.

3. Dolce, G., et al., Care and neurorehabilitation in the disorder of consciousness: a model in progress. ScientificWorldJournal, 2015. 2015: p. 463829.

4. Katz, D.I., et al., Natural history of recovery from brain injury after prolonged disorders of consciousness: outcome of patients admitted to inpatient rehabilitation with 1-4 year follow-up. Prog Brain Res, 2009. 177: p. 73–88.

5. Giacino, J.T., K. Kalmar, and J. Whyte, The JFK Coma Recovery Scale-Revised: measurement characteristics and diagnostic utility. Arch Phys Med Rehabil, 2004. 85(12): p. 2020–9.

6. American Congress of Rehabilitation Medicine, B.I.-I.S.I.G.D.o.C.T.F., et al., Assessment scales for disorders of consciousness: evidence-based recommendations for clinical practice and research. Arch Phys Med Rehabil, 2010. 91(12): p. 1795–813.

7. Schnakers, C., et al., Diagnostic accuracy of the vegetative and minimally conscious state: clinical consensus versus standardized neurobehavioral assessment. BMC Neurol, 2009. 9: p. 35.

8. Tamashiro, M., et al., A Spanish validation of the Coma Recovery Scale-Revised (CRS-R). Brain Inj, 2014. 28(13-14): p. 1744–7.

9. Schnakers, C., et al., A French validation study of the Coma Recovery Scale-Revised (CRS-R). Brain Inj, 2008. 22(10): p. 786–92.

10. Sacco, S., et al., Validation of the Italian version of the Coma Recovery Scale-Revised (CRS-R). Brain Inj, 2011. 25(5): p. 488–95.

11. Zaytsev, О., The Psychopathology of Severe Brain Injure. Book in Russian ed. 2011, Moscow: MEDPRESS-inform. 336.

12. Bykova, V.I., V.I. Lukyanov, and E.V. Fufaeva, Dialogue with the patient in low consciousness state after severe brain damages. Counseling Psychology and Psychotherapy. 23(3): p. 9–31.

13. Avezaat, C.J., R. Braakman, and A.I. Maas, [A scoring device for the level of consciousness: the Glasgow "coma" scale]. Ned Tijdschr Geneeskd, 1977. 121(53): p. 2117–21.

14. Giacino, J.T., et al., Monitoring rate of recovery to predict outcome in minimally responsive patients. Arch Phys Med Rehabil, 1991. 72(11): p. 897–901.

15. Wijdicks, E.F., et al., Validation of a new coma scale: The FOUR score. Ann Neurol, 2005. 58(4): p. 585–93.

16. Kevric, J., et al., Validation of the Full Outline of Unresponsiveness (FOUR) Scale for conscious state in the emergency department: comparison against the Glasgow Coma Scale. Emerg Med J, 2011. 28(6): p. 486–90.

17. Fischer, M., et al., Inter-rater reliability of the Full Outline of UnResponsiveness score and the Glasgow Coma Scale in critically ill patients: a prospective observational study. Crit Care, 2010. 14(2): p. R64.

18. Wannez, S., et al., Prevalence of coma-recovery scale-revised signs of consciousness in patients in minimally conscious state. Neuropsychol Rehabil, 2017: p. 1–10.

19. Schnakers, C., et al., Does the FOUR score correctly diagnose the vegetative and minimally conscious states? Ann Neurol, 2006. 60(6): p. 744–5; author reply 745.

20. Squires, J.E., et al., Validation of the conceptual research utilization scale: an application of the standards for educational and psychological testing in healthcare. BMC Health Serv Res, 2011. 11: p. 107.

21. Hinkle, D.E., W. Wiersma, and S.G. Jurs, Applied statistics for the behavioral sciences. 5th ed ed. 2003, Boston: Houghton Mifflin.

22. Marcati, E., et al., Validation of the Italian version of a new coma scale: the FOUR score. Intern Emerg Med, 2012. 7(2): p. 145–52.

23. Neese, L.E., et al., Neuropsychological assessment and the Disability Rating Scale (DRS): a concurrent validity study. Brain Inj, 2000. 14(8): p. 719–24.

24. Shiel, A., et al., The Wessex Head Injury Matrix (WHIM) main scale: a preliminary report on a scale to assess and monitor patient recovery after severe head injury. Clin Rehabil, 2000. 14(4): p. 408–16.

25. La Porta, F., et al., Can we scientifically and reliably measure the level of consciousness in vegetative and minimally conscious States? Rasch analysis of the coma recovery scale-revised. Arch Phys Med Rehabil, 2013. 94(3): p. 527–535 e1.

26. Lovstad, M., et al., Reliability and diagnostic characteristics of the JFK coma recovery scale-revised: exploring the influence of rater’s level of experience. J Head Trauma Rehabil, 2010. 25(5): p. 349–56.

